# A Molecular Signature for Anastasis, Recovery from the Brink of Apoptotic Cell Death

**DOI:** 10.1101/102640

**Authors:** Gongping Sun, Elmer Guzman, Hongjun Robin Zhou, Kenneth S. Kosik, Denise J. Montell

## Abstract

During apoptosis, executioner caspase activity has been considered a point of no return. However, recent studies show that cells can survive caspase activation following transient apoptotic stimuli, a process named anastasis. To identify a molecular signature, we performed whole transcriptome RNA sequencing of untreated, apoptotic, and recovering HeLa cells. We found that anastasis is an active, two-stage program. During the early stage, cells transition from growth-arrested to growing. In the late stage, cells change from proliferating to migratory. Strikingly, some early recovery mRNAs were elevated first during apoptosis, implying that dying cells poise to recover, even while still under apoptotic stress. Furthermore, TGFβ-induced Snail expression is required for anastasis, and recovering cells exhibit prolonged elevation of pro-angiogenic factors. This study demonstrates similarities in the anastasis genes, pathways, and cell behaviors to those activated in wound healing. This study identifies a repertoire of potential targets for therapeutic manipulation of this process.

## Introduction

Apoptosis is a cell suicide program that is conserved in multicellular organisms and functions to remove excess or damaged cells during development and stress^1,2^. Excessive apoptosis contributes to degenerative diseases, whereas blocking apoptosis can cause cancer^3^. Apoptotic cells exhibit distinctive morphological changes^4^ caused by activation of proteases called caspases^5,6^. Activation of executioner caspases is a necessary step during apoptosis^5^ and until recently was considered a point of no return^7^.

However executioner caspase activation is not always sufficient to kill cells under apoptotic stress. For example, caspase 3 activation in cells treated with sub-lethal doses of radiation or chemicals does not cause morphological changes or death, but rather allows cells to survive with caspase-dependent DNA damage that can result in oncogenic transformation^8–10^. In addition, transient treatment of cells with lethal doses of certain apoptosis inducers causes caspase 3 activation sufficient to cause apoptotic morphological changes, yet cells can survive after removing the toxin in a process called anastasis^11^. While most cells fully recover, a small fraction bear mutations, and an even smaller fraction undergo oncogenic transformation. Cell survival following executioner caspase activation has also been reported in cardiac myocytes responding to transient ischemia, in neurons over-expressing Tau, and during normal Drosophila development^12–15^. Taken together these studies suggest that cells can recover from the brink of apoptotic cell death and that this can salvage cells, limiting the permanent tissue damage that might otherwise be caused by a transient injury. However, the same process of anastasis in cancer cells might underlie recurrence following transient chemo- or radiation therapy. Thus, defining the molecular changes occurring in cells undergoing this remarkable recovery from the brink of death is a critical step toward manipulating this survival mechanism for therapeutic benefit.

## Results

### RNAseq reveals anastasis composes of two stages

To initiate apoptosis, we exposed HeLa cells to a 3 hr treatment with ethanol (EtOH), which was sufficient to induce cell shrinkage and membrane blebbing (Figure 1A, B), cleavage of PARP1, which is a target of caspase 3/7 (Figure 1I), and activation of a fluorescent reporter of caspase 3 activity, in ~75% of the cells (Figure G, H, J, supplementary video 1). Removal of the EtOH by washing allowed a striking recovery to take place over the course of several hours, during which time ~70% of the cells re-attached to the culture matrix and spread out again (Figure 1C-F, K, supplementary video 2)^11^.

**Figure 1.**
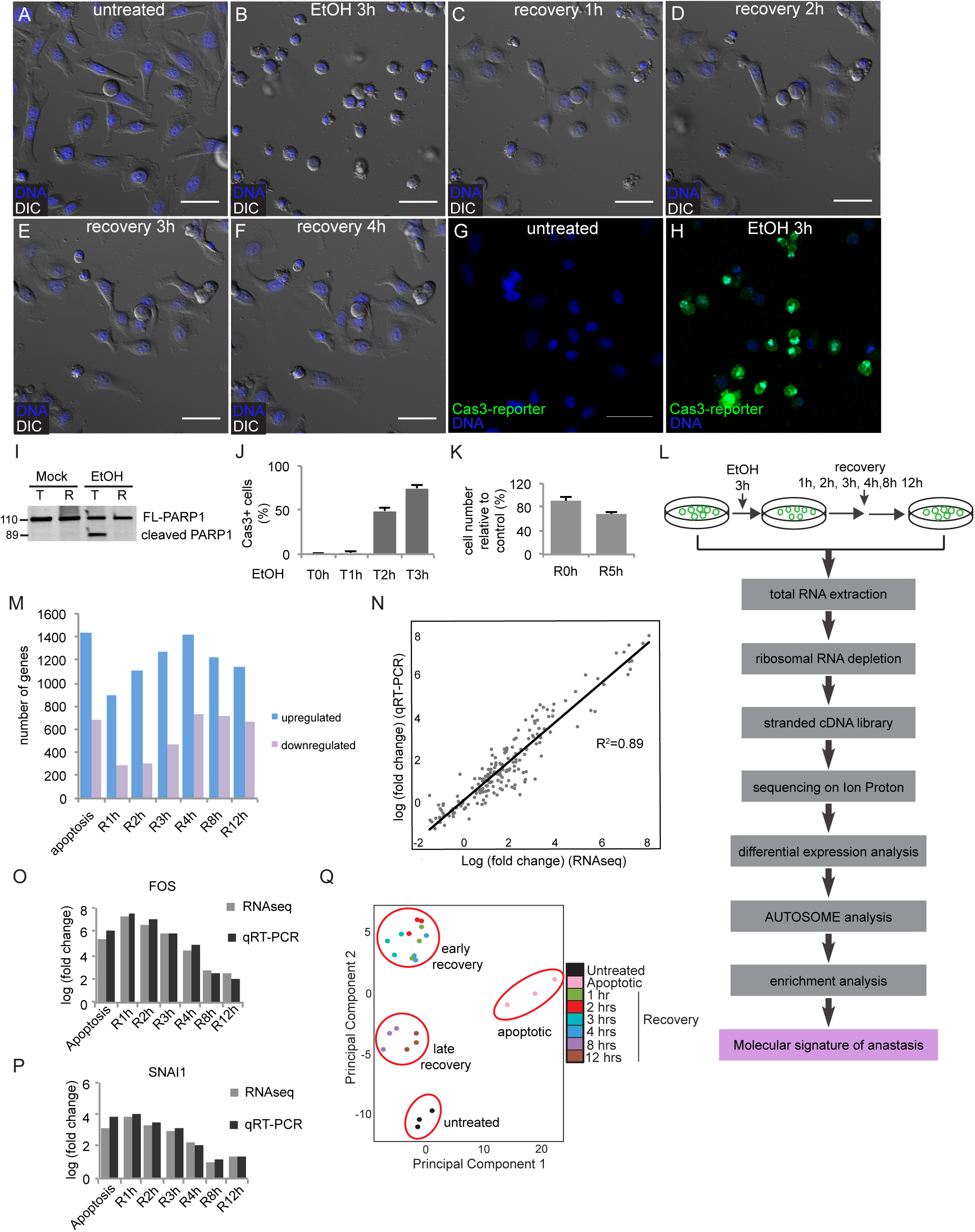
**RNAseq defines anastasis as a two-stage active process**. A-F) Time-lapse live imaging of HeLa cells before EtOH treatment (A), after 3hr EtOH treatment (B), and after recovery for 1hr (C), 2hrs (D), 3hrs (E), and 4hrs (F). G-H) Caspase 3 activity (green fluorescence) in the same group of cells before (G) and after 3hr EtOH treatment. DAPI staining is shown in blue in A-H. Scale bar is 50μm. I) Western blots of full-length PARP1 (FL-PARP1) and cleaved PARP1 in cells after 3hrs mock or EtOH treatment (T) followed by 21hrs recovery (R). J) Quantification of the percentage of cells with active caspase 3 during EtOH treatment (n=5). K) The ratio of the number of remaining cells right after washing away EtOH (R0h) or after 5hr recovery (R5h) to the number of cells after mock treatment (n=3). L) The workflow of RNAseq experiments. M) Numbers of upregulated and downregulated genes in apoptotic cells, cells after 1hr, 2hr, 3hr, 4hr, 8hr, and 12hr recovery compared to untreated cells (fold change>1.5, false discovery rate < 0.05). N) Correlation of RNAseq and qRT-PCR data for 27 genes. O, P) Comparison of the changes in levels of FOS (O) and SNAI1 (P) mRNAs over time detected by RNAseq and qRT-PCR. In n-p, ‘fold change’ is compared to the expression level in untreated cells. Q) Principal component analysis of RNAseq data reveals four clusters: untreated cells, apoptotic cells, 1-4 hrs recovery, and 8 and 12 hrs recovery. Each color represents a different time point. Each time point was analyzed in triplicate.

To define anastasis at a molecular level, we performed whole transcriptome RNA sequencing (RNAseq) of untreated cells, apoptotic cells, and of cells allowed to recover for 1, 2, 3, 4, 8, or 12 hours (Figure 1L). These time points include and extend beyond the time needed for the major morphological changes, which appeared complete by 4 hours (Figure 1A-F). Compared to untreated cells, 900-1500 genes increased in abundance >1.5-fold at each time point, while 250-750 genes decreased >1.5 fold (false discovery rate <0.05) (Figure 1M, Supplementary table S1). Well-characterized genes such as Fos, Jun, Klf4, and Snail were induced, as well as genes about which little is known, such as the long non-coding RNA LOC284454, a gene predicted to encode a deubiquitinating enzyme (OTUD1), a pseudokinase (TRIB1), and a phosphate carrier protein (SLC34A3).

We validated the expression patterns of 27 top-ranked differentially expressed (22 upregulated and 5 downregulated) genes using quantitative reverse transcription PCR (qRT-PCR). The results of RNAseq and qRT-PCR correlated well, with an R^2^ of 0.89 (Figure 1N-P, Supplementary figure S1). Out of 22 top-ranked, up-regulated genes we tested, 18 increased in abundance in HeLa cells recovering from apoptosis induced by a second chemical (10% DMSO), and in a second cell line (human glioma H4 cells) recovering from EtOH-induced apoptosis (Supplementary figure S2). This suggests that there is a core anastasis response that is neither cell-line-specific nor apoptosis-inducer-specific.

A principle component analysis (PCA) of the RNAseq data showed that cells undergoing anastasis clustered into two distinct groups: one group composed of cells allowed to recover for 1-4 hours, and a second containing cells that recovered for 8 or 12 hours. Both groups were also clearly different from apoptotic cells and untreated cells (Figure 1Q). We therefore defined the first 4 hrs of recovery as the early stage, and 8 to 12 hrs as late.

### Distinct features of early and late recovery

To compare the transcriptional profiles between early and late stages, we used the program AUTOSOME^16^, which clusters genes according to similarities in their expression patterns over time (Supplementary table S2). This approach identified 8 clusters containing a total of 1,172 genes upregulated during early recovery, and 6 clusters containing 759 genes upregulated late (Supplementary table S3). We refer to these as early and late response genes, respectively (Figure 2A, Supplementary table S3). Gene Ontology (GO) analysis revealed enrichment of expected categories such as “regulation of cell death” and “cellular response to stress” in the early response (Figure 2B). The GO term “transcription” was the most significantly enriched, indicating induction of transcription factors during initiation of anastasis (Figure 2B). The term “chromatin modification” was also enriched. Enrichment of early response genes in “regulation of cell proliferation” and “regulation of cell cycle” terms suggested that removing apoptotic stress released cells from a growth-arrested state to re-enter the cell cycle and proliferate (Figure 2B). Remarkably, the classes of early and late response genes were very different. The late response was enriched in posttranscriptional activities such as ncRNA processing and ribosome biogenesis.

**Figure 2.**
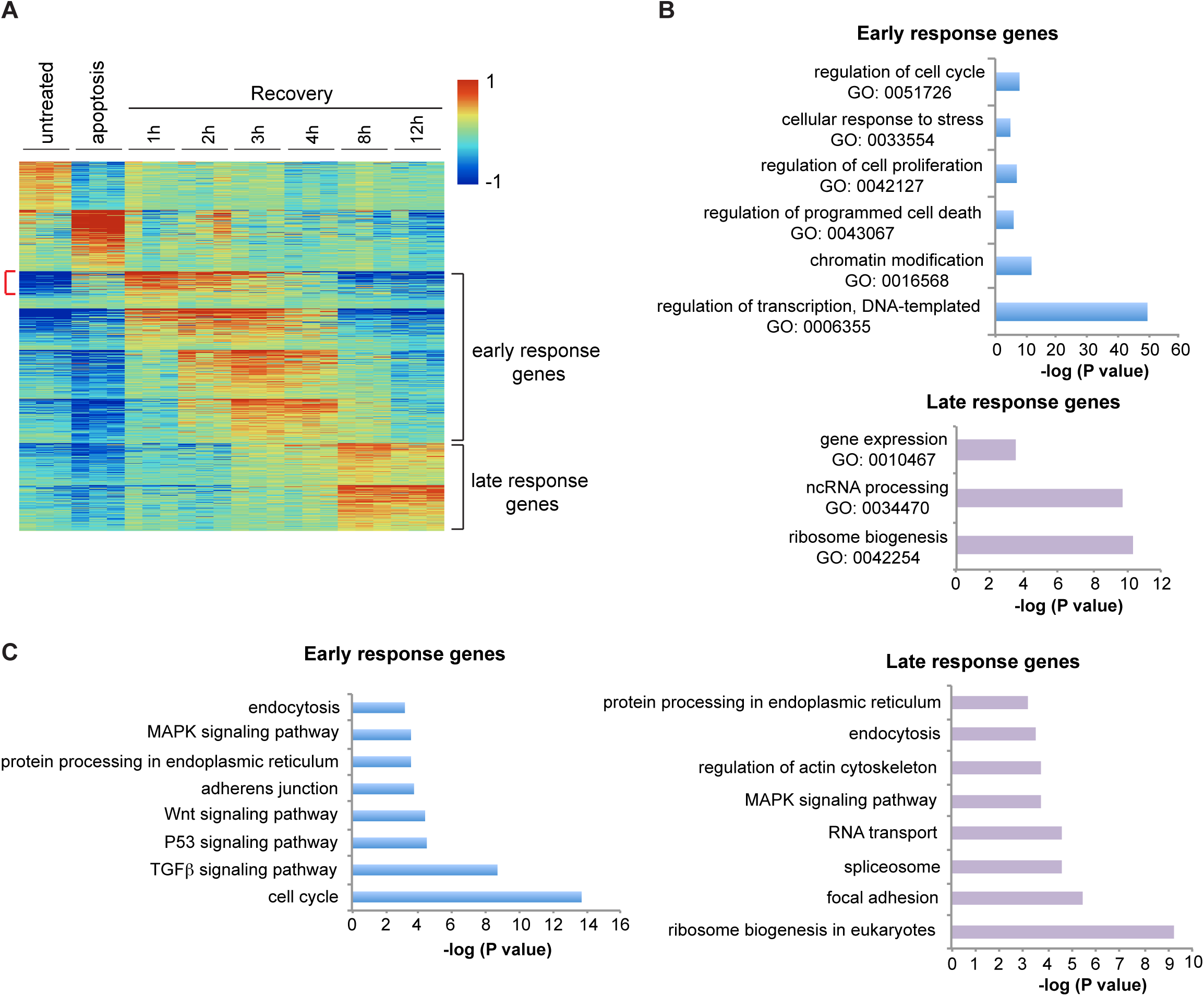
**AUTOSOME and enrichment analyses of early and late response genes**. A) AUTOSOME analysis. The red indicates increases and blue indicates decreases in mRNA abundance. Genes most highly upregulated during 1 to 4hrs recovery are defined as early response genes, and those that peak at 8 or 12hrs are late response genes. Red bracket points out the genes upregulated in both apoptosis and early recovery. B) GO enrichment analysis of early response genes and late response genes. C) KEGG pathway enrichment analysis of early and late response genes. In B and C), P value is the Bonferroni P value.

KEGG pathway analysis showed early response genes to be enriched in cell cycle and pro-survival pathways such as TGFβ, MAPK, and Wnt signaling (Figure 2C). The late response showed enrichment in general post-transcriptional pathways such as ribosome biogenesis, RNA transport, protein processing, and endocytosis, as well as specific pathways such as focal adhesion and regulation of actin cytoskeleton (Figure 2C).

### Cells transition from proliferation to migration during recovery from apoptosis

Consistent with enrichment of cell cycle and proliferation genes in the early response, cell numbers increased during the first 11 hours of recovery (Figure 3A, Supplementary figure S3), plateaued after 11 hours, and began to increase again at ~ 30 hours. At even later time points (after replating), recovered cells exhibited a similar proliferation rate to control, mock-treated cells (Figure 3B).

**Figure 3.**
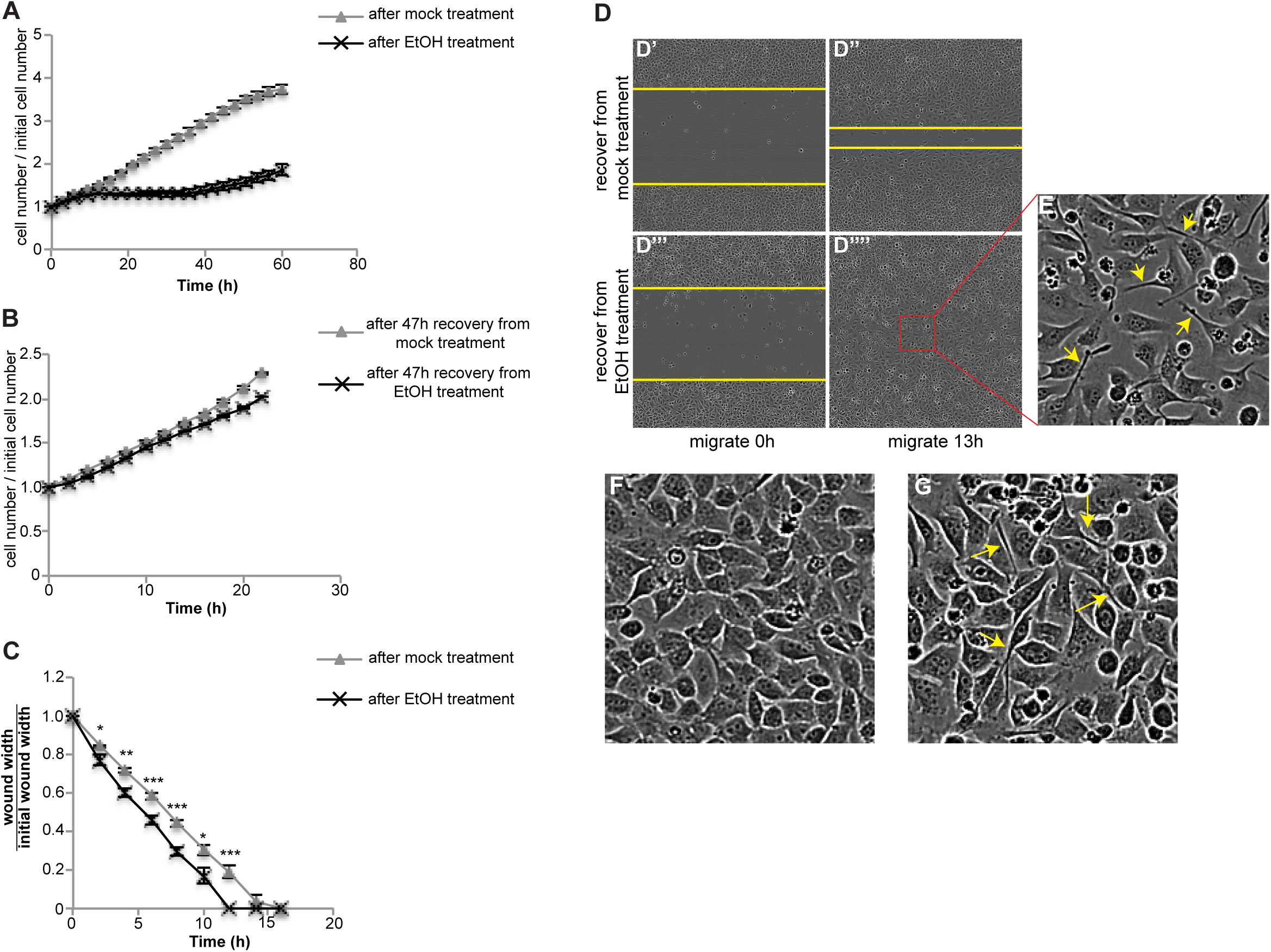
**Cells transition from proliferation to migration during recovery**. A) Cell number during recovery after mock or EtOH treatment (n=3). B) Cells after 47hr recovery from mock or EtOH treatment were trypsinized and re-plated at similar density. The curves show the change of cell number over 22hrs (n=3). C, D) wound healing assays. C) Quantification of wound width over time (n=8). The stars above each time point present the statistical significance of the difference between cells after mock versus EtOH treatment. *: P<0.05. **: P<0.01. ***: P<0.001. ****: P<0.0001. D) Images of wounds made in cells recovering from mock treatment (D’ and D’’) or EtOH treatment (D’’’ and D’’’’). The yellow lines mark the wound margins. E) Magnified images of the outlined regions in (D’’’’). F and G) Images of confluent monolayers of cells recover from mock (F) or EtOH (G) treatment. In E and G, yellow arrows point to elongated cells. In all plots, error bars represent standard error of mean.

Due to the enrichment of “focal adhesion” and “regulation of actin cytoskeletion” pathways in late response gene clusters (Figure 2C), we hypothesized that cells might become migratory during the proliferation pause. To measure migration, we performed wound-healing assays. Scratch wounds made in monolayers of cells allowed to recover from EtOH treatment for 16 hrs closed faster than that in mock-treated monolayers (Figure 3C, D), even though they exhibited a lower cell number and slightly slower proliferation rate (Supplementary figure S4). In both mock-treated and EtOH-treated cells, those that migrated to fill the wound were more elongated than cells lagging behind (Figure 3E). A larger proportion of cells recovering from EtOH treatment showed this elongated morphology compared to mock-treated cells (Figure 3F, G), suggesting this morphology might facilitate migration and wound closure.

### Recovery from apoptotic stress is different from recovery from autophagy

The observed enrichment of cell cycle components in the early response suggested that one facet of anastasis is re-entry into the cell cycle following growth arrest during apoptosis. To distinguish which molecular features of anastasis were common to another type of growth arrest and recovery, and to identify those more likely to be specific to anastasis, we evaluated the expression of the top-ranked, differentially expressed anastasis genes in cells undergoing recovery from nutrient deprivation. Nutrient deprivation induces growth arrest and autophagy, a process that can promote survival^17,18^. Autophagy results in degradation of cytoplasmic components in autophagosomes, which are double-membrane-bound vesicles that sequester cytoplasm and fuse with lysosomes^17^. However expression of autophagy genes was not induced during anastasis, suggesting that the two survival mechanisms differ. A time course showed that amino acid starvation for 2 hours induced autophagy in HeLa cells, shown by increased LC3 staining, which is a marker of for autophagosomes. LC3 staining is typically further augmented by blocking fusion between autophagsomes and lysosomes with bafilomycin A1^19^, and this was also true in nutrient-deprived HeLa cells (Figure 4A-D). Two hours of starvation did not induce caspase 3 activation (Figure 4E), although longer treatments did. Of the 24 genes upregulated during anastasis that we tested, 10 were downregulated or only slightly upregulated during recovery from autophagy (Figure 4F-O). Thus elevated transcription of these 10 genes distinguishes cells in early anastasis from those recovering from autophagy. Furthermore cells recovering from autophagy showed no measurable difference in the rate of wound closure compared to mock treated cells (Figure 4P, Q). Thus, cells recovering from transient apoptotic stress exhibit both molecular and behavioral hallmarks that distinguish anastasis from recovery from other types of stress that induce growth arrest.

**Figure 4.**
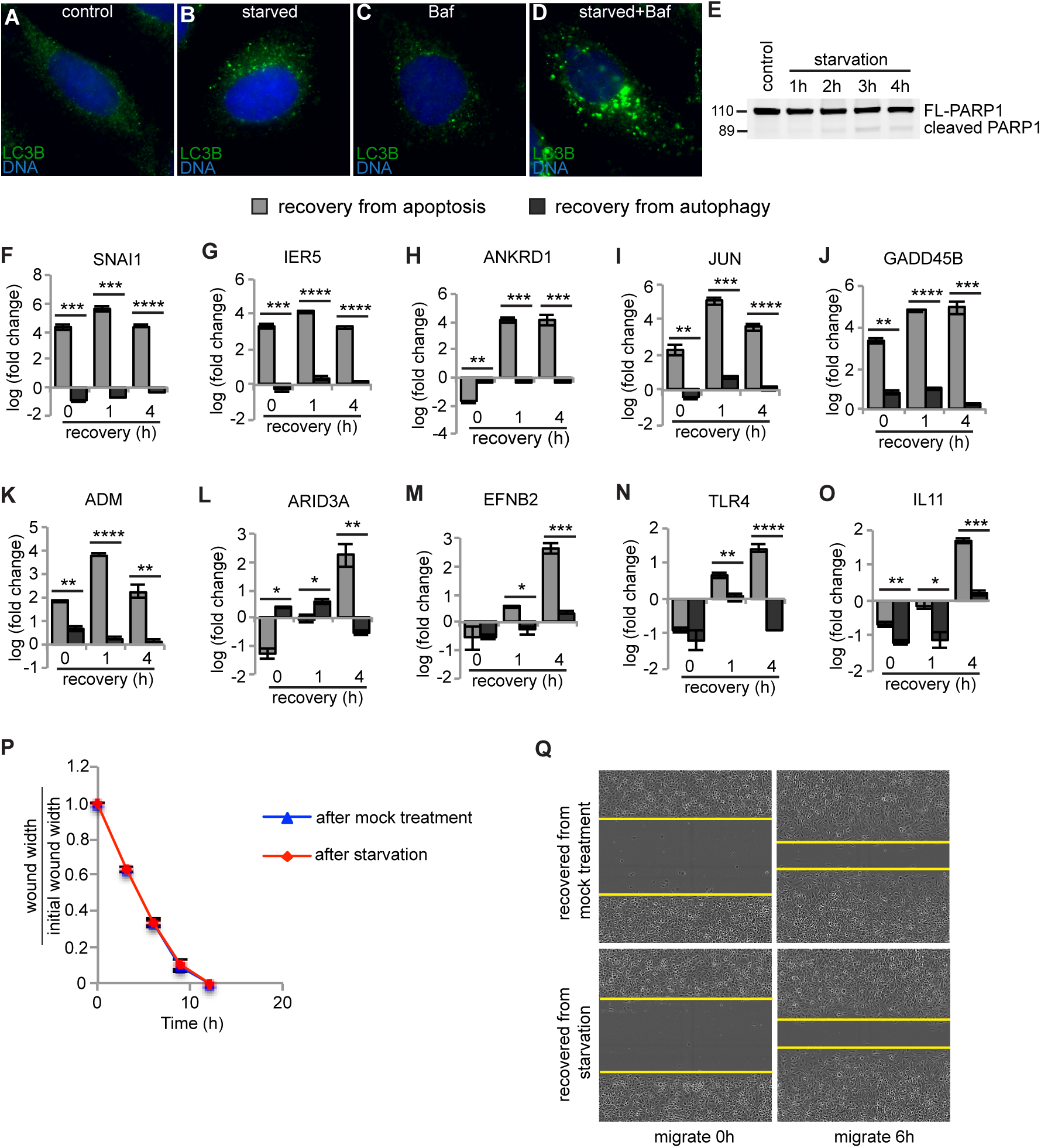
**Comparison between anastasis and recovery from autophagy** A-D) LC3B autophagosome marker staining (green) in cells incubated with growth medium containing 1% DMSO (control) (A), or HBSS containing 1% DMSO (B), or growth medium containing 100nM Bafilomycin A1 (C), or HBSS containing Bafilomycin A1 (D). E) Western blot for PARP1 showing little cleavage during amino acid starvation. The 2hr time point was chosen for further studies. F-O) Comparison between the mRNA levels of indicated genes after 0hr, 1hr, and 4hr recovery from apoptosis (gray bars) or from autophagy (black bars). ‘fold change’ is compared to expression level of mock-treated cells. *: P<0.05. **: P<0.01. ***: P<0.001. ****: P<0.0001. n=3. P) Relative wound width over time in wound healing assay (n=5). Q) Images of wounds made in cells recovering from mock treatment (upper two panels) or amino acid starvation (lower two panels). Yellow lines mark the margin of wound. In all bar graphs, error bars represent standard error of mean.

### Cells poise for recovery during apoptosis

The transcriptional profile of cells undergoing anastasis revealed an unknown feature of apoptotic cells that appears to contribute the rapid transition to recovery. We noticed that transcripts corresponding to a subset of early response genes that were induced during the first hour of anastasis were already elevated in abundance in apoptotic cells, relative to untreated cells (Figure 1O, P, Figure 2A, Supplementary 1A-G). One possible explanation would be that these are genes that drive apoptosis, and that apoptosis had not completely stopped 1 hour after removal of the chemical stress. Alternatively, these could be genes encoding proteins that contribute to recovery, and that cells prepare for that possibility even after caspase activation. To distinguish between these opposing possibilities, we compared the levels of expression of 10 such early genes at 1 hr of recovery after 3 hr EtOH treatment, to the levels in cells left in EtOH for 4 hrs. The mRNA levels after 4 hr EtOH treatment were significantly lower than those at 1 hr recovery, indicating that accumulation of these mRNAs was associated with the survival response (Figure 5A-J). We analyzed the corresponding protein levels for five of the early response targets for which antibodies were available. The protein levels remained unchanged or were slightly reduced during apoptosis (Figure 5K), suggesting that while the mRNAs accumulated, their translation was inhibited, consistent with prior observations of downregulated protein synthesis in apoptotic cells^20^. This intriguing finding supports the idea that even during apoptosis, cells actually poise for recovery by synthesizing, or protecting from degradation, specific mRNAs encoding survival proteins, which are however not translated. If apoptotic stress persists, the mRNAs are degraded and the cells die. However, if the apoptotic stress disappears, cells are prepared to rapidly synthesize survival proteins. This ‘poised for recovery’ state may help to explain the rapid recovery after stress removal.

**Figure 5.**
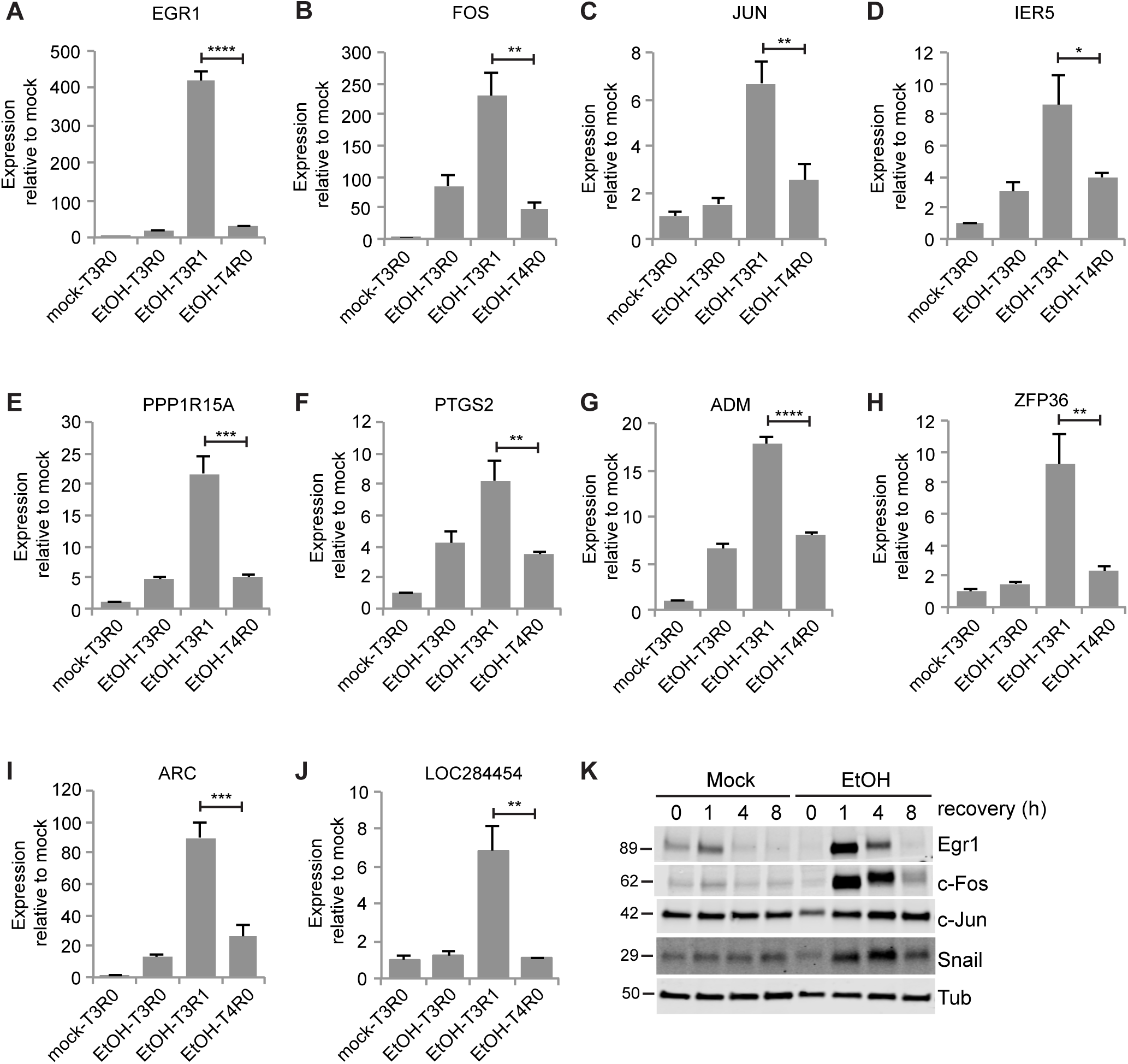
**Apoptotic cells were poised for recovery**. A-J) mRNA levels of the indicated genes relative to mock-treated cells after 3hr mock treatment (mock-T3R0), after 3hr EtOH treatment (EtOH-T3R0), after 1hr recovery from EtOH treatment (EtOH-T3R1) and after 4hr EtOH treatment (EtOH-T4R0) (n=3). Stars show statistic significance between EtOH-T3R1 and EtOH-T4R0. *: P<0.05. **: P<0.01. ***: P<0.001. ****: P<0.0001. K) The protein levels of Egr1, c-Fos, c-Jun, Snail during recovery from mock treatment and EtOH treatment.

### Knocking down Snail impaired recovery from apoptosis

Snail is one of the mRNAs enriched in apoptotic cells and then highly induced in early recovery, and has been reported to protect cells from apoptosis^
21,22^. We found that Snail protein levels increased during recovery (Figure 5K). In addition, knocking down Snail expression by stably expressing short hairpin RNA (shRNA) reduced the endogenous Snail protein level (Figure 6A) and reduced the percentage of cells surviving EtOH-induced apoptosis (Figure 6B). We also found increased PARP1 cleavage in Snail-depleted cells following EtOH treatment (Figure 6A). Therefore the poor recovery following Snail knockdown may result from enhanced caspase activation during EtOH treatment, impaired anastasis, or both.

**Figure 6.**
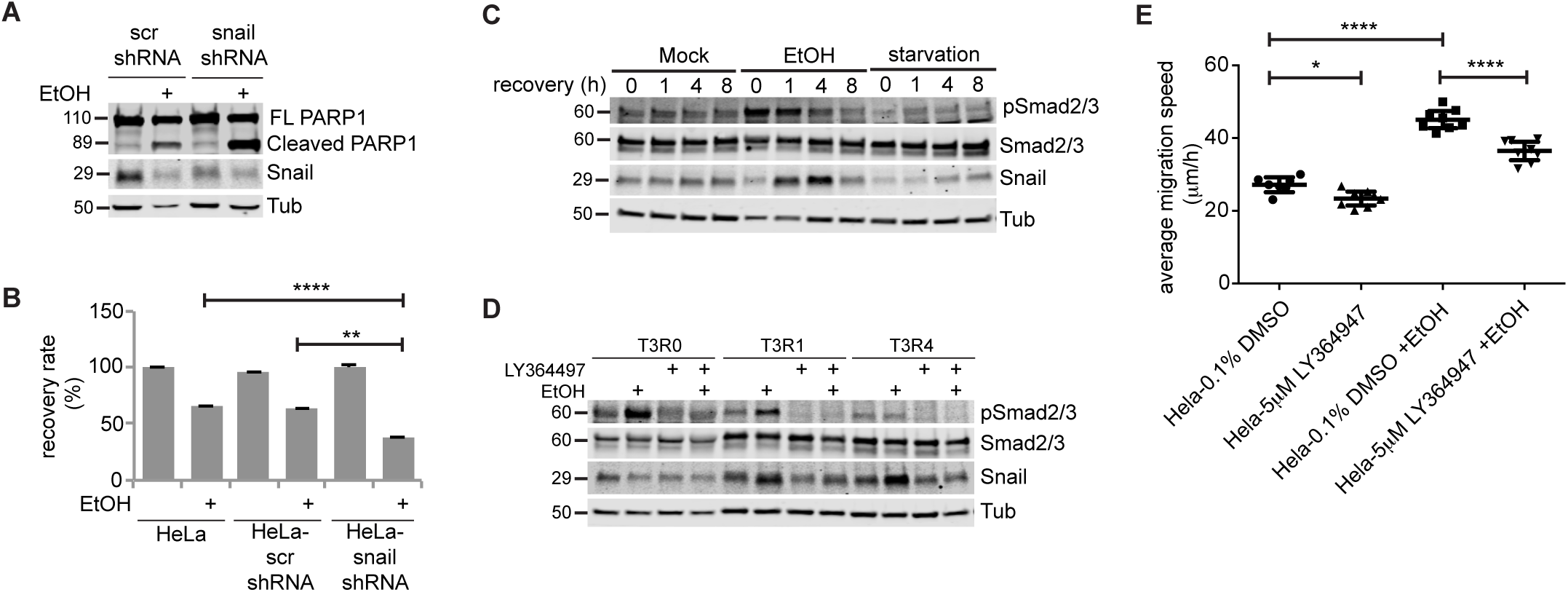
**Transient activation of TGFβ signaling induced Snail upregulation, which was required for anastasis, and caused increased migration in late stage**. A) Cleavage of PARP1 and Snail protein in HeLa cells stably expressing scrambled shRNA (scr shRNA) or snail shRNA treated with or without EtOH for 3hrs. B) Recovery rate of untransfected HeLa, HeLa-scr shRNA, and HeLa-snail shRNA cells after mock treatment or EtOH treatment (n=3). In all bar graphs, error bars represent standard error of mean. C) Western blots of phosphor-Smad2/3, total Smad2/3, Snail in cells recovering from mock treatment, EtOH treatment or starvation. D) The level of pSmad2/3, Smad2/3 and Snail in apoptotic cells (T3R0), cells after 1hr recovery (T3R1) and 4hr recovery (T3R4). The addition of LY364497 and EtOH is indicated. E) Average migration speed of the indicated group of cells during wound healing assay (n=8). Before wound healing assay cells were treated with or without EtOH together with 0.1% DMSO or 5μM LY364947 for 3 hrs, followed by 4 hrs recovery with 0.1% DMSO or 5μM LY364947 and an additional 16 hrs recovery without any inhibitor. Error bars represent 95% confidence interval. *: P<0.05. **: P<0.01. ***: P<0.001. ****: P<0.0001.

### Activation of TGFβ signaling contributes to Snail upregulation and migration

One important upstream regulator of Snail is TGFβ signaling^23^, and this pathway was enriched in the early recovery gene set. TGFβ signaling regulates transcription through phosphorylation and activation of downstream transcription factors Smad2 and Smad3^24^. Phosphorylation of Smad2/3 increased during apoptosis and in the first hour of recovery, then diminished after 4 hrs of recovery, indicating transient activation of TGFβ signaling (Figure 6C). To test if Snail upregulation was due to TGFβ signaling, we treated cells with a TGFβ receptor I specific inhibitor LY364947. LY364947 did not affect basal cell survival or proliferation (Supplementary figure S5) but suppressed the induction of Snail protein during recovery (Figure 6D).

TGFβ signaling activation can promote epithelial-to-mesenchymal transition (EMT) and cell migration^25^. To determine if the transient activation of TGFβ signaling during early recovery was responsible for the increased migration later, we inhibited TGFβ signaling only during apoptosis and early recovery stage, then tested cell migration using the wound-healing assay. Transient inhibition of TGFβ signaling by LY364947 reduced the average migration speed of EtOH-treated cells from 45 to 37 μm/h while reducing the average migration speed of mock-treated cells from 27 to 23 μm/h, suggesting TGFβ signaling contributes to both basal motility and anastasis-induced migration in HeLa cells (Figure 6E). Interestingly, TGFβ signaling, Snail mRNA, and Snail protein were all downregulated during autophagy and recovery (Figure 4F, Figure 6C). Recovery from autophagy did not stimulate cell migration (Fig 4P, Q), further supporting the idea that activation of TGFβ signaling, induction of Snail, and increased migration characterize the recovery from the brink of apoptotic cell death specifically, rather than general stress responses.

### Induction of angiogenesis-related genes throughout recovery

While TGFβ signaling and Snail expression were transiently elevated during early recovery, some angiogenesis-related genes were persistently elevated throughout the 12 hours examined. Placenta growth factor (PGF) binds vascular endothelial growth factor receptor (VEGFR) and stimulates endothelial cell proliferation and migration^26^. PGF was among the top 10 upregulated genes at every time point during apoptosis and recovery (Supplementary table S1). PGF mRNA increased ~22-fold at 1hr recovery, and even after 24hr recovery, PGF mRNA was 3-fold higher in EtOH-treated cells compared to mock-treated cells (Figure 7A, B).

**Figure 7.**
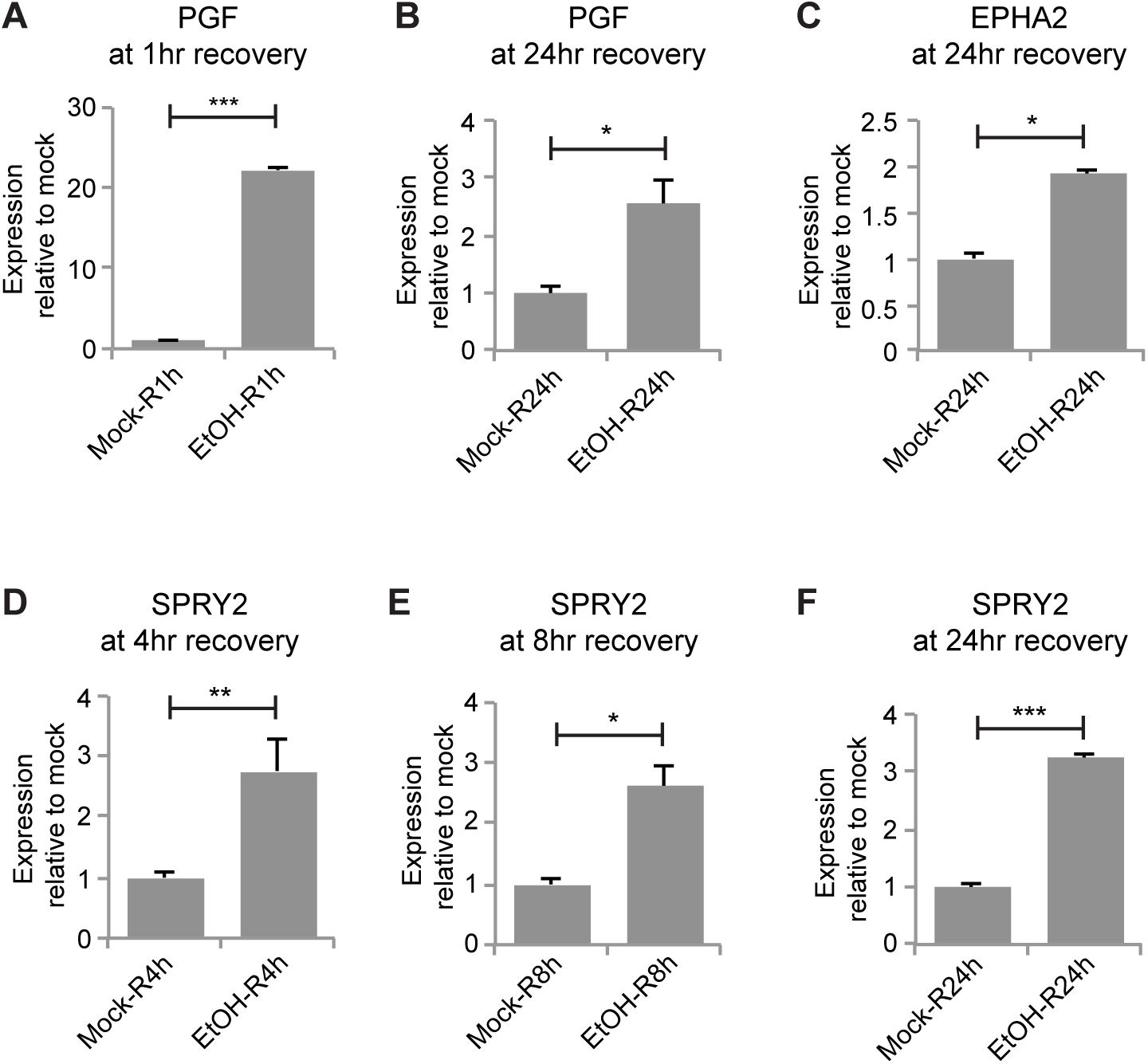
**Angiogenesis-related genes were persistently upregulated during recovery**. mRNA expression of PGF (A, B), EPHA2 (C), SPRY2 (D-F) in cells recover from mock or EtOH treatment for the indicated time (R). Error bars represent standard error of mean. *: P<0.05. **: P<0.01. ***: P<0.001. ****: P<0.0001.

Ephrin and Ephrin receptor (EphR) signaling are also important in blood vessel development and angiogenesis ^27,28^. Several EphRs (EPHA2, EPHB2, EPHB4) and Ephrins (EFNB1, EFNB2) were upregulated throughout recovery (Supplementary table S1). For example, expression of EFNB2 in the first hour of recovery was ~1.6 fold that of mock-treated cells and elevated ~ 2-3 fold during 3-12 hours of recovery (Supplementary figure S1K). EPHA2 was significantly upregulated after 24 hr recovery (Figure 7C). Sprouty 2 (SPRY2) is a common transcriptional target of VEGFR and EphR signaling^29^. SPRY2 expression was upregulated from 4hr to 24hr recovery (Figure 7D-F), suggesting activation of VEGFR and EphR signaling during recovery.

## Discussion

The ability of cells to survive caspase-3 activity has implications for normal development, cancer, and degenerative and ischemic diseases. Here we report the first molecular characterization of cells recovering from the brink of apoptotic cell death. The data show that anastasis proceeds in two clearly defined stages that are characterized by distinct repertoires of genes. In the early stage, cells transcribe mRNAs encoding many transcription factors and re-enter cell cycle. In the late stage, cells pause in proliferation while increasing migration. While the proliferation and migration responses were transient, others were longer lasting. For example, we found that cells that have undergone anastasis elevate expression of angiogenesis-related genes for 24 hours. In vivo, these factors would be expected to exert a non-autonomous effect of stimulating blood vessel growth. Taken together the results presented here demonstrate that cells recovering from the brink of apoptotic cell death express factors that promote proliferation, survival, migration, and angiogenesis (Figure 8). The cell biological processes involved in anastasis are thus reminiscent of wound healing responses ^30^, consistent with the idea that cells evolved this capacity in order to limit permanent tissue damage following a transient injury.

**Figure 8.**
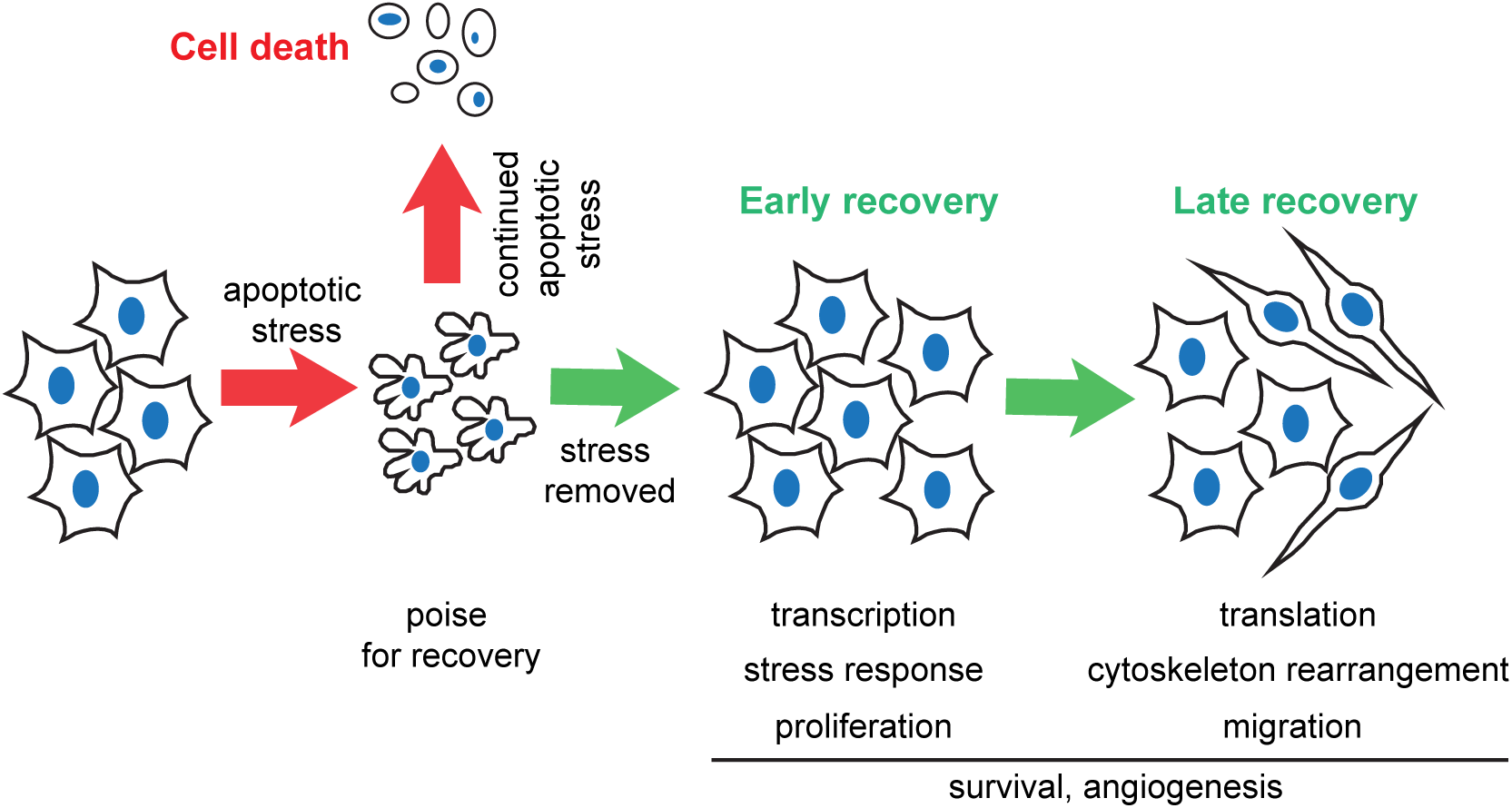
**Schematic of events in anastasis**. During apoptosis, cells poise for recovery. If the stress persists, cells die. If the stress is removed, cells undergo two stages of recovery.

Many of the same molecular pathways are also upregulated during wound healing and in cells undergoing anastasis including TGFβ, receptor tyrosine kinase, MAPK signaling, and angiogenesis promoting pathways^30^. TGFβ signaling and Snail expression are thought to promote EMT and chemotherapy resistance during tumor progression^31–33^. Cells undergoing anastasis activate TGFβ signaling and Snail expression, and become migratory, all features of EMT. Two recent studies reported that EMT, though dispensable for tumor metastasis, is required for tumor recurrence after chemotherapy^34,35^, suggesting that EMT is a survival strategy for tumor cells under stress *in vivo*. Our results suggest a possible relationship between tumor recurrence, EMT, and anastasis. If cancer cells exposed to radiation or chemotherapy during treatment escape death via anastasis, TGFβ signaling and Snail expression would be induced, and these critical regulators of EMT^32^ would confer resistance to further chemotherapy and radiation^33^. Therefore anastasis could in principle drive EMT-dependent tumor recurrence.

In vivo, fast growing tumors require formation of new blood vessels to supply nutrients and provide a route to metastasis^36^. Common cancer treatments, like tumor resection, can induce extensive angiogenesis, which may promote tumor recurrence^37^. In fact, elevated expression of angiogenic factors and/or increased blood vessel formation have been found in recurrent craniopharyngioma, bladder cancer, squamous cell carcinoma after surgery or irradiation^38–40^. Our results imply if tumor cells survive apoptosis triggered by chemotherapy, irradiation or surgery, these survivors may upregulate production of pro-angiogenic factors to facilitate angiogenesis and tumor recurrence.

In addition to providing insight into the molecular nature of anastasis, the work presented here uncovers an unanticipated aspect of apoptosis. Even during apoptosis, cells poise for recovery by accumulating mRNAs encoding survival proteins. The proteins however are not expressed until and unless they are needed to reverse the apoptotic process. We propose that this facilitates rapid recovery upon removal of the apoptotic stimulus. Apoptosis and caspase 3 have been linked to tumor recurrence. In cancer patients, the rate of recurrence is positively correlated with the amount of activated caspase 3 in tumor tissues^41,42^. One explanation that has been offered to explain this somewhat paradoxical result is that after radiotherapy or chemotherapy, activated caspase 3 promotes production of pro-growth signals that are released from dying cells to stimulate proliferation of living tumor cells, leading to tumor recurrence^41,43,44^. Dying cells can also secrete VEGF in a caspase-dependent way to promote angiogenesis after irradiation^45^. These previous studies proposed that apoptotic cells induce non-autonomous compensatory proliferation in neighboring cells. The model is that cells that activate caspase 3 die and stimulate cells without activated caspase to proliferate and grow. However, our study suggests another possible mechanism underlying tumor recurrence: that tumor cells with activated caspase 3 themselves may eventually survive, proliferate, migrate, and trigger angiogenesis, contributing to tumor repopulation.

When facing tissue injury, anastasis could in principle facilitate repair and regeneration, and limit the permanent damage that might otherwise occur in response to a powerful but temporary insult. On the other hand, anastasis would be detrimental if adopted by cancer cells in response to chemo or radiation therapy, thus potentially promoting recurrence. Thus the mechanisms described here fit into the general idea that cancers mimic and coopt wound-healing behaviors^46^. Enhancing anastasis would be expected to be beneficial in the context of degenerative or ischemic disease, whereas inhibiting anastasis should be beneficial in cancer treatment.

## Materials and methods

### Cell culture

Human cervical cancer HeLa cells (ATCC cell line CCL-2), was grown in MEM supplemented with GlutaMAX (Thermo Fisher Scientific), 10% fetal bovine serum (Sigma), and 100U/ml Penicillin-Streptomycin (Thermo Fisher Scientific). Human neuroglioma H4 cells (ATCC cell line HTB-148) were grown in high glucose DMEM supplemented with GlutaMAX (Thermo Fisher Scientific), 10% fetal bovine serum, and 100U/ml Penicillin-Streptomycin. All cells were maintained at 37°C with 5% CO_2_ and 90% humidity. Cells were tested for Mycoplasma contamination.

For RNAseq, 1.2×10^6^ cells were seeded in each 100mm dish and cultured overnight. The next day, cells were treated with either fresh growth medium or fresh growth medium with 4.3% ethanol (EtOH) (Fisher Scientific) for 3 hrs. Samples from untreated and apoptotic cells were collected at this moment. For recovery, medium was carefully removed and fresh growth medium was added. For each time point, three biological replicates were included.

For qRT-PCR, western blotting, and immunofluorescent antibody staining, 2×10^5^ cells were seeded in a 35mm dish or 6-well plate and cultured overnight. The next day, cells were treated with either fresh growth medium, or fresh growth medium with 4.3% (HeLa) or 4% (H4) EtOH for 3 hrs or fresh growth medium with 10% dimethyl sulfoxide (DMSO) (Santa Cruz Biotechnologies) for 2.5 hrs, or Hanks’ balanced salt solution (HBSS) (Thermo Fisher Scientific) for 2 hrs. The precise concentration and time points were chosen based on titration studies to achieve the highest possible percentage of apoptotic cells that could recover. For recovery, medium was carefully removed and fresh growth medium was added for the indicated period of time.

For TGFβ signaling inhibition, 5μM LY364947 (Sigma) was added to cells together with mock or EtOH treatment.

### RNA extraction

For RNA sequencing, total RNA was extracted using mirVana miRNA isolation kit (Thermo Fisher Scientific) then treated with TURBO Dnase (Thermo Fisher Scientific) to get rid of the genomic DNA. Ribosomal RNA (rRNA) was removed using RiboMinus Eukaryote System v2 (Thermo Fisher Scientific). The quality of RNA was examined using fragment analyzer (Advanced Analytical).

For qRT-PCR, RNA was extracted using RNeasy Mini kit (Qiagen) and treated with TURBO DNase to remove genomic DNA.

### RNA sequencing and data analysis

cDNA libraries used for sequencing were made from rRNA-depleted RNA using Ion Total RNA-seq kit v2 (Thermo Fisher Scientific), and sequenced on an Ion Torrent Proton sequencer (Thermo Fisher Scientific). Strand specific single-end reads were generated from sequencing with average read lengths of 75 bp. Reads were mapped to UCSC Human Reference Genome (hg19) using Tophat (v2.0.13)^47^. Reads covering gene coding regions were counted using htseq (v0.6.1)^48^ and resulting count data was used for was used for downstream analysis. Count data were first filtered by removing genes with low expression, or genes with less than 50 reads in more than 2 replicates per sample. Remaining count data was normalized using trimmed mean of M-values method using edgeR (v 3.14.0)^49^. Normalized count distributions were fit to a generalized linear model in order to test for differential expression of genes (p-value=0.05) among multiple samples. The differential expression test was corrected for multiple testing by applying the Benjamini-Hochberg method on p-values to control false discovery rate. AutoSOME^16^ was used for identification of gene clusters with similar expression patterns on CPM (counts per million) and log2 transformed count data.

Gene Ontology enrichment analysis was performed using DAVID (https://david.ncifcrf.gov/) and PANTHER (http://pantherdb.org). Only the common GO terms with Bonferroni P value less than 0.001 and FDR less than 0.001 were considered significantly enriched. KEGG pathway enrichment analysis was performed using WebGestalt (http://www.webgestalt.org).

### qRT-PCR

RNA samples were reverse transcribed into cDNA using SuperScript III first-strand synthesis system (Thermo Fisher Scientific). And qPCR was performed on QuantStudio 12K Flex real-time PCR system (Thermo Fisher Scientific) with Power SYBR green PCR master mix. The primers used in qPCR are listed in supplementary table S4.

### Live imaging

2×10^5^ cells were seeded in each glass-bottom, 35mm dish (MatTek corporation). Cells were incubated with Hoechst 33342 (Molecular Probes) for 20min, and imaged on Zeiss LSM 780 with temperature and CO_2_ control. Images were taken every 10min. Medium change for EtOH treatment and recovery was carried out between scans. To monitor caspase 3 activation during EtOH treatment lively, NucView 488 (Biotium) was added. NucView488 binds irreversibly to DNA and thus inhibits anastasis.

### Western blotting

Cells were lysed in laemmli sample buffer and run in 4-20% Mini-PROTEAN TGX precast protein gels (Bio-rad). Primary antibodies used were rabbit anti-Egr1 (Cell signaling #4154), rabbit anti-c-Fos (Cell signaling #2250), rabbit anti-c-Jun (Cell signaling #9165), mouse anti-Snail (Cell signaling #3895), rabbit anti-PARP1 (Cell signaling #9532), rabbit anti-Smad2/3 (Cell signaling #8685), rabbit anti-pSmad2/3 (Cell signaling #8828), mouse anti-α-Tubulin (Sigma #T6199). Secondary antibodies used were IRDye 800CW donkey anti-rabbit IgG (H+L), IRDye 680LT donkey anti-mouse IgG (H+L), IRDye 800CW donkey anti-mouse IgG (H+L) (Li-Cor Biosciences). The blots were scanned on Odyssey imaging system (Li-Cor Biosciences). Cell treatment, sample collection and western blotting were repeated at least three times, and the representative blots were shown in the figures.

### Immunofluorescent staining

Cells were seeded in 6-well plate with coverslip. After treatment, cells were washed once with PBS and fixed with cold methanol for 5min. Cells were then rinsed with PBS twice and washed with PBS containing 0.2% Triton X-100 (Fisher Scientific). After that, cells were blocked with PBS containing 0.2% Triton X-100 and 5% goat serum (Sigma). The primary antibody and secondary antibody used are rabbit anti-LC3B (Cell signaling #2775) and Alexa Fluor 488 conjugated goat anti-rabbit IgG (H+L) secondary antibody (Thermo Fisher Scientific). The images were acquired on a Leica DMi8 microscope. Cell treatment and staining were repeated three times, and the representative images were shown in the figure.

### Short-hairpin RNA construct, transfection and stable cell line

Short-hairpin RNA constructs were made in the pLVX vector. The sequences of snail shRNA and scrambled shRNA were 5’-GGATCTCCAGGCTCGAAAGtcaagagCTTTCGAGCCTGGAGATCCtttttt-3’^50^ and 5’-CCTAAGGTTAAGTCGCCCTCGCTCGAGCGAGGGCGACTTAACCTTAGGttttt-3’ (Addgene #1864). The constructs were transfected into HeLa cells using TurboFect transfection reagent (Thermo Fisher Scientific), and selected using 2μg/ml puromycin (Thermo Fisher Scientific) to get stable cell lines.

### Proliferation assay

HeLa NucLight Red cells (Essen BioScience) were seeded in 6-well plate and cultured overnight. After EtOH treatment, cells were cultured in growth medium and imaged in the IncuCyte Zoom (Essen BioScience) every hour. 9 fields of view were taken per well, and the number of red fluorescent nuclei was counted using IncuCyte Zoom software (Essen BioScience).

### Wound healing assay

HeLa NucLight Red cells were seeded in 100mm dish and cultured overnight. After treatment and 16hr recovery, cells were trypsinized and seeded in Matrigel-coated 96- well ImageLock plate (Essen BioScience) at 4×10^4^ cells per well. After 4hrs, wound was made in each well using WoundMaker (Essen BioScience). The wound closure process was imaged in IncuCyte Zoom every hour.

### Recovery rate quantification

Before treatment, cells seeded in 6-well plate were incubated with growth medium with DRAQ5 (Thermo Fisher Scientific) for 10min at 37C, and imaged in IncuCyte Zoom to quantify the original cell number. After 4hr recovery, cells were washed twice with PBS to remove floating dead cells, and stained with DRAQ5 again. The cell number was quantified again as the number of survivors. The recovery rate was calculated as the ratio between the number of survivors to the original cell number.

### Statistical analyses

Statistical analysis used in RNAseq data analysis was described in the “RNA sequencing and data analysis” section. For other experiments, statistical significance was determined using unpaired, two-tailed, t-tests with Welch’s correction for comparison between two samples, and one-way ANOVA to compare more than two samples, with P<0.05 set as criteria for significance. The Tukey test was used to derive adjusted P value for pairwise comparison among multiple samples. Sample size was not predetermined.

### Data availability

The RNAseq data have been deposited in the Gene Expression Omnibus (GEO) under accession ID GSE86480.

## Acknowledgements

This work was supported by NIH grant NIH DP1OD019313-01 and NIH 5DP1CA195760-02 to Denise J. Montell. We thank Ugochukwu Ihenacho, Jing Xiong, Rebecca Cheng, Kirsten Wong for technical assistance.

### Author contributions

E. G. participated in preparing libraries for RNAseq and carried out bioinformatics analyses of the data. H. R. Z. carried out the mapping of RNAseq reads. K. K. provided advice on the design, execution, and interpretation of the RNAseq data. G. S. carried out the majority of experiments presented and participated in preparing libraries for RNAseq and in the bioinformatics analysis. G. S. also prepared the figures and wrote the first draft of the manuscript. D.J.M. conceived of and coordinated the study, advised G. S. in all aspects of the work, contributed to design and interpretation of the results, and revised the manuscript.

### Conflict of interest

The authors declare no conflict of interest.

